# Nuclear Tau modulates VGluT1 expression: a new function for Tau

**DOI:** 10.1101/394312

**Authors:** Giacomo Siano, Martina Varisco, Maria Claudia Caiazza, Valentina Quercioli, Marco Mainardi, Chiara Ippolito, Antonino Cattaneo, Cristina Di Primio

## Abstract

**Summary**

Tau displacement from microtubules is the first step in the onset of tauopathies, and is followed by toxic protein aggregation. However, other non-canonical functions of Tau might have a role in these pathologies. Here, we demonstrate that a small amount of Tau localizes in the nuclear compartment and accumulates in both the soluble and DNA-bound fractions. We show that nuclear Tau regulates the expression of VGluT1, a disease-relevant gene directly involved in glutamatergic synaptic transmission. Impeding Tau/tubulin interaction in the cytosol favours its nuclear translocation and increases VGluT1 expression. Remarkably, the P301L mutation impairs this mechanism leading to a loss of function. Altogether, our results provide the demonstration of a direct physiological role of Tau on gene expression. Alterations of this mechanism may be at the basis of the onset of neurodegeneration.

## Introduction

The main known physiological function of Tau is to bind and stabilize microtubules (MTs) [1–3]. This interaction influences MTs stability, axonal transport and growth [3–6] and its alteration has a relevant role in tauopathies. During the progression of the pathology, Tau undergoes post-translational modifications, leading to detachment from MTs and an ensuing increase of the soluble pool of Tau that is considered more easily subjected to post translational modifications and truncation. This event is thought to determine aggregation and the progression of neurodegeneration, with deficits in synaptic transmission, neuronal loss and cognitive impairment [1,2,7,8].

Despite the high and increasing incidence of these diseases, currently available drugs do not halt their progression, pointing up the need to understand the molecular mechanisms involved in early steps of tauopathy.

All current approaches to target Tau are based on its known actions on MTs, and are aimed at counteracting the aggregation of soluble pool of Tau to block its seeding and spreading. Non canonical functions of Tau, not linked to its microtubule-binding properties, have been suggested [9–16], but their relevance in physiology and pathology is not known. A small amount of Tau has been found to be localized within the nucleus and a growing body of evidence supports the affinity of Tau for nucleic acids. *In vitro* studies demonstrated that Tau is able to bind the minor groove of dsDNA [17,18] and this interaction takes place with both AT- and GC-rich oligonucleotides [17,19]. Tau has been found, in neuroblastoma and neuronal nuclei [9,13,20], associated with heterochromatin and pericentromeric DNA [14,16,21], but Tau function in this compartment is still unclear. A putative role in nucleolar organization has been suggested [9,13,14]. Moreover, a possible role of Tau in DNA damage protection and chromosome stability has been proposed, since wild-type Tau prevents DNA damage under oxidative stress or hyperthermic conditions [10,16,18] while Tau bearing the pathological mutation P301L is associated to chromosome instability and aneuploidy [22–24].

Nevertheless, although these observations suggest a role of Tau in the nuclear compartment, this field is still largely unexplored and it is not known whether nuclear Tau could have a role in modulating gene expression or some aspects of chromatin function that might be relevant for the pathophysiology of tauopathies.

Here, we provide the first demonstration that the nuclear pool of Tau modulates the expression of the vesicular glutamate transporter VGluT1, which is known to be strongly upregulated in early phases of tauopathies [25,26]. Moreover, we show that increasing the soluble pool of Tau favours its nuclear translocation and increases the expression of VGluT1. The Tau^P^301^L^ mutation hinders this chain of events and impairs its effect on VGluT1 expression.

Our results provide new information on the physiological role of Tau and, above all, suggest important implications for the pathogenesis of tauopathies.

## Results

### Tau is detectable in the soluble and chromatin-bound nuclear fractions, and increases VGluT1 mRNA and protein levels

To study the possible role of nuclear Tau in chromatin functions, we performed a sub-cellular fractionation and isolated the soluble and DNA-bound nuclear fractions of the hippocampal neuronal cell line HT22 and the neuroblastoma cell line SHSY5y as established models to study the neuronal cytotoxicity in neurodegeneration. Beside the endogenous expression of Tau proteins, we overexpressed the wild-type Tau isoform 4R0N.

We verified that Tau is clearly detectable in the soluble nuclear fraction; moreover, we showed that it accumulates in the DNA-bound fraction (Figure 1A), confirming in a cellular context previous evidence on Tau-DNA interaction *in vitro* [17,27].

**Figure 1.**
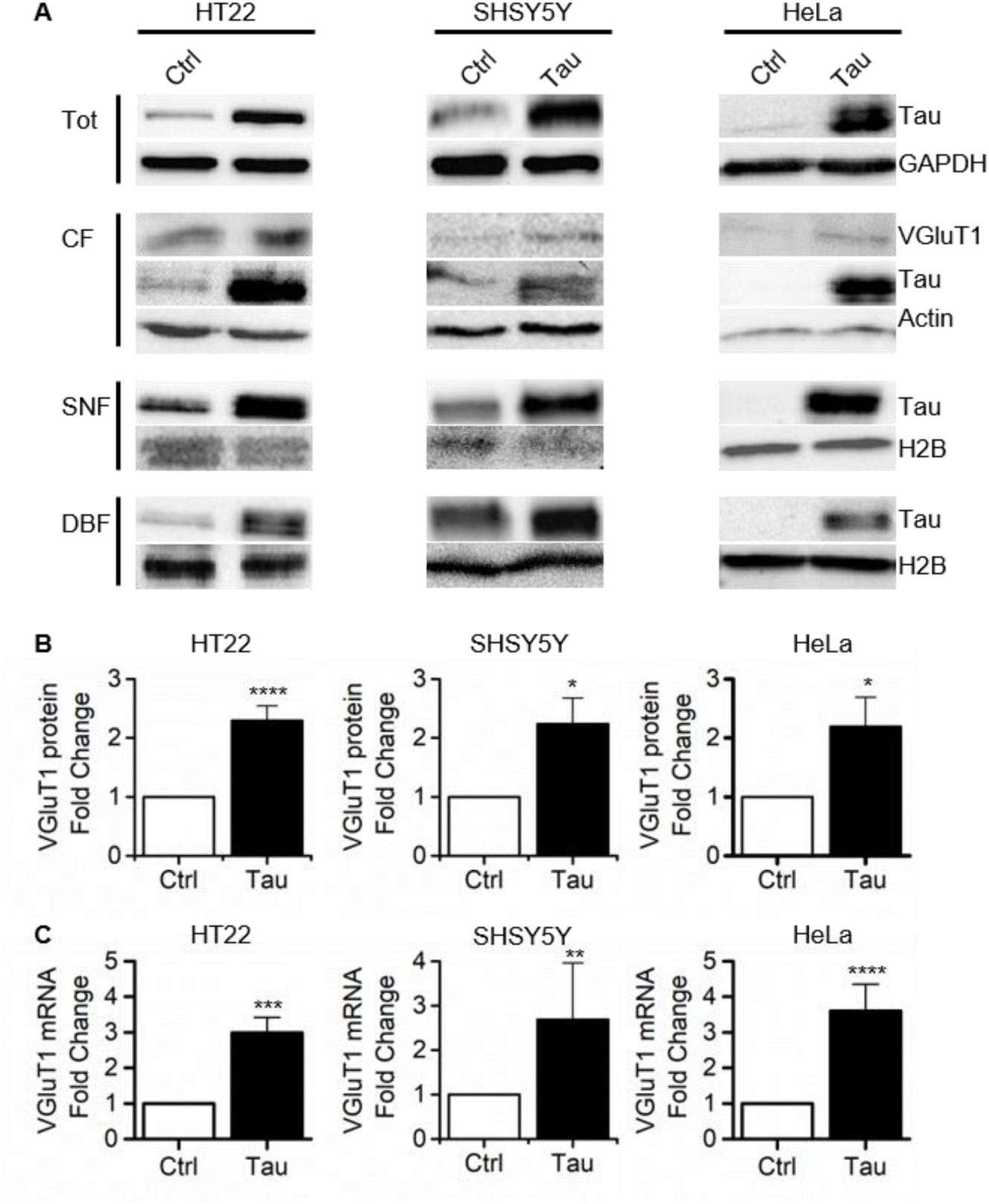
Tau is detectable in nuclear fractions and increases VGluT1 expression A. Western blot of subcellular fractions obtained from neuronal (HT22 and SHSY5Y) and non-neuronal (HeLa) cell lines. **B**. Quantification of VGluT1 protein level in control and Tau-expressing cells **C**. VGluT1 mRNA quantification by qPCR. CF: cytoplasmic fraction; SNF: soluble nuclear fraction; DBF: DNA-bound fraction. (Mann-Whitney test; **** p<0.0001; *** p<0.001) (See also Figure S1).

To investigate whether nuclear Tau might have a role in modulating gene expression, we checked the expression of disease-related genes in Tau-overexpressing cells. A growing body of evidence suggests that glutamate release is altered during the progression of tauopathies, causing an early synaptic hyperexcitability in the asymptomatic phase mediated by Tau [11,25,26].

Therefore, we focused on the presynaptic vesicular glutamate transporters (VGluT1) that package glutamate into vesicles for neurotransmitter exocytosis [28].

We observed an increase of VGluT1 protein level (HT22: 2.2±0.26SE Fold Change=FC; SHSY5Y: 2.24±0.44SE FC) in neuronal cell lines overexpressing Tau (Figure 1A-B). Remarkably, we also found that Tau expression increases VGluT1 mRNA (HT22: 2.99±0.43SE FC; SHSY5Y: 2.69±1.27SE FC) (Figure 1C).

To verify whether this effect is determined by the interference of the overexpressed Tau with the endogenous isoforms we performed the same analysis on non-neuronal cells (Hela cells) that do not express endogenous Tau. We found that in this cellular background the de novo expression of Tau induced the increased expression of VGluT1 mRNA and protein as well (mRNA: 3.6±0.75SE FC; protein: 2.19±0.49SE FC) (Figure 1).

Taken together, these results indicate that Tau modulates VGluT1 gene expression, in a cell type-independent manner.

### VGluT1 increased expression is dependent on nuclear Tau

To investigate whether the modulation of VGluT1 expression could be mediated by the nuclear pool of Tau or by an indirect effect of cytoplasmic Tau, we forced the localization of Tau in the nucleus or its retention in the cytoplasm by including a nuclear localization sequence (Tau-NLS) or a nuclear export signal (Tau-NES), respectively. As expected, Tau-NLS prevalently accumulated in the nuclear fraction, while Tau-NES was mainly detected in the cytoplasm (Figure 2A, Figure S1).

**Figure 2.**
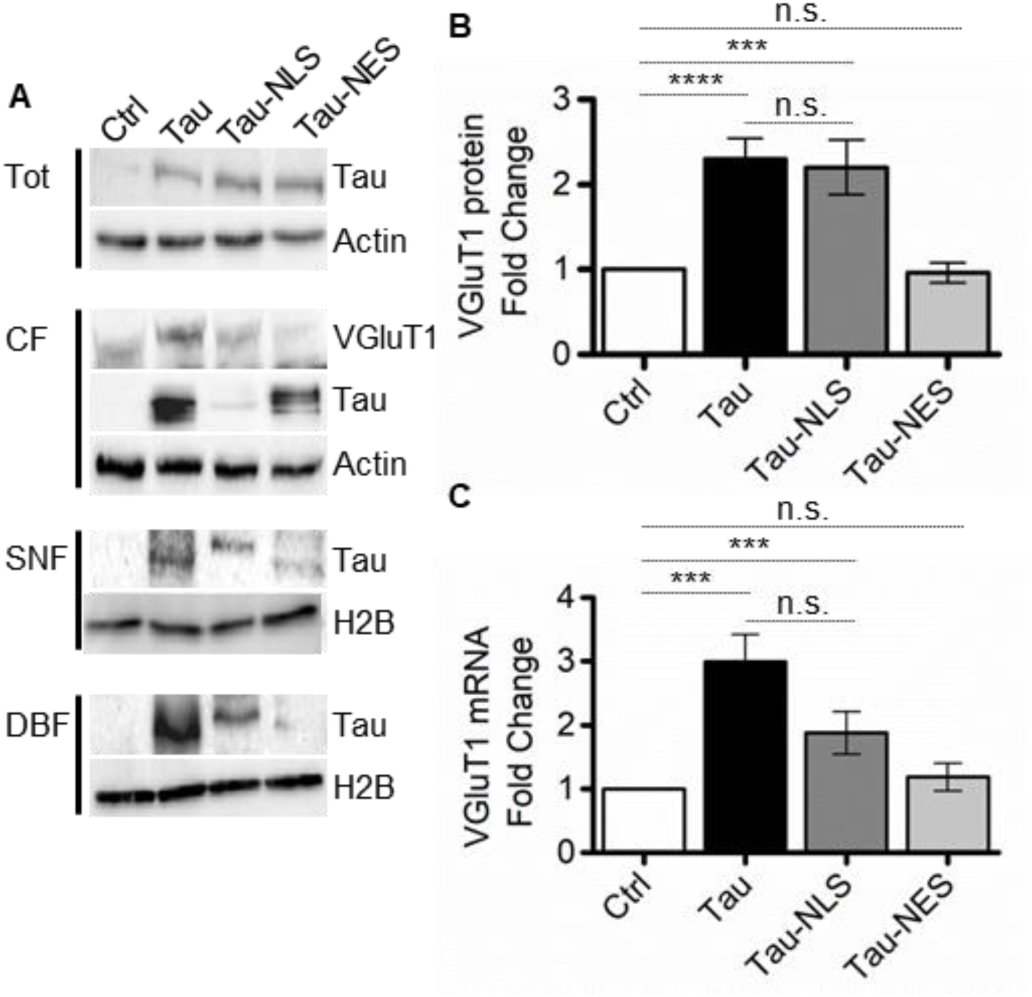
VGluT1 increased expression is mainly dependent on nuclear Tau A. Western blot of HT22 cells expressing Tau-NLS or Tau-NES. (See also Figure S2) **B**. Quantification of WB experiments. **C**. VGluT1 mRNA quantification by qPCR in HT22 cells. (Mann-Whitney test; **** p<0.0001; *** p<0.001; n.s. p>0.05).

We observed that Tau-NLS induced an increase in VGluT1 protein expression (2.0±0.36SE FC) comparable to that induced by untagged Tau. On the contrary, when the nuclear localization of Tau was prevented (Tau-NES), no increase in VGluT1 protein could be observed (Figure 2A-B).

Likewise, cells expressing Tau-NES showed no increase in the level of VGluT1 mRNA, while cells expressing Tau-NLS showed an increase similar to that observed with untagged Tau (Tau-NLS: 1.88±0.33SE FC; Tau-NES: 1.18±0.21SE FC) (Figure 2C).

This set of data indicates that Tau proteins translocated into the nucleus regulate the expression of VGluT1. Accordingly, shifting the equilibrium towards cytoplasmic Tau, by forcing its nuclear export, brings VGluT1 expression back to baseline.

### Displacing Tau from microtubules increases the localization of endogenous Tau in the nucleus

At the onset of pathology, Tau undergoes post-translational modifications that induce its detachment from microtubules. The increased availability of soluble Tau protein is currently believed to slowly favour the toxic oligomerization [1,2,29,30]. We reasoned that this event could shift the equilibrium between the Tau pools in the cytoplasmic and nuclear compartments, thus increasing Tau nuclear localization. To investigate this hypothesis, we induced the detachment of endogenous Tau from MTs by treating HT22 cells with paclitaxel (PTX) or nocodazole (Noc). These drugs increase soluble Tau with two different mechanisms, preserving or disrupting the MT network, respectively. Indeed, PTX competes with Tau for the same binding pocket on the tubulin dimer, causing its displacement from MTs; nocodazole disassembles the MT network, leading to an increase in soluble Tau [31–34]. Importantly, to avoid possible misleading results due to Tau overexpression, we treated neuron-like cells expressing the endogenous but not the exogenous Tau. As showed by WB experiments, in treated cells we observed an increase of Tau levels in both the nuclear soluble fraction and in DNA-bound nuclear fraction (Figure 3A, Figure S2). We conclude that the destabilization of the endogenous Tau from MTs favours not only an increase in the cytoplasmic soluble pool, but also its translocation to the nuclear compartment.

**Figure 3.**
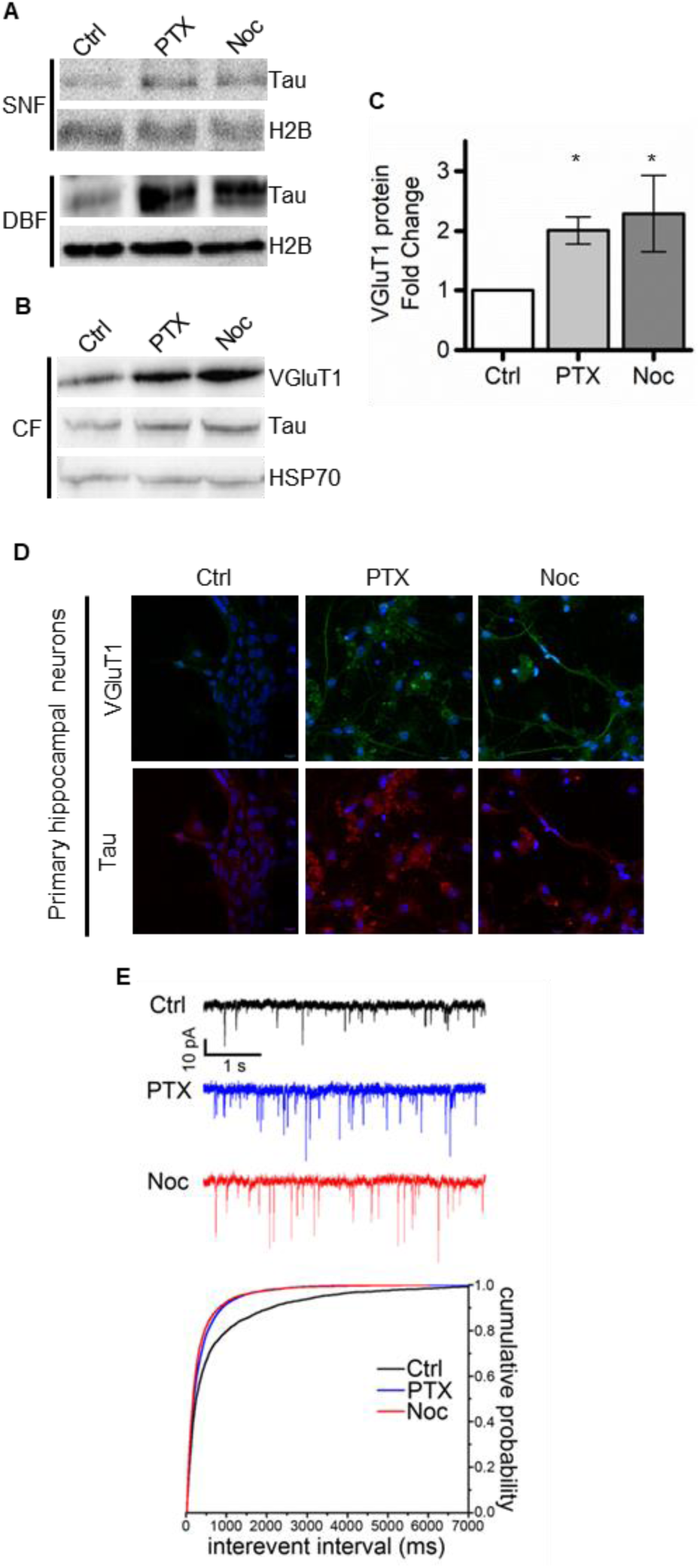
Tau displacement from MTs increases Tau nuclear accumulation and VGluT1 expression A-B. Western blot analysis of subcellular fractions from untreated and treated cells. UT (untreated), PTX (paclitaxel), Noc (nocodazole). **C**. Quantification of VGlut1 protein level (B) (Mann-Whitney test; * p<0.05; see also Figure S3) **D**. Representative confocal imaging of IF in primary neurons. Tau (red), VGluT1 (green), DAPI (blue). **E.** Cumulative distributions of mEPSC frequency (i.e., interevent interval) following either PTX or Noc treatment, compared to untreated controls (Ctrl, n = 3002 events from 12 cells; PTX, n = 4911 events from 10 cells; Noc, n = 3576 events from 6 cells; Kolmogorov-Smirnov test, Ctrl vs. PTX, p<0.001; Ctrl vs. Noc, p<0.001; PTX vs. Noc, p<0.001).

### Displacing Tau from microtubules increases the expression of VGluT1 and glutamatergic synaptic transmission

Having established that the amount of nuclear Tau can be directly influenced by modulating its interaction with cytoplasmic MTs, we investigated whether increasing the nuclear pool of endogenous Tau determines a concomitant increase in VGluT1 expression. We observed that in PTX- and Noc-treated cells VGluT1 expression rises significantly (PTX, 2.0±0.23SE FC; Noc, 2.29±0.64SE FC) (Figure 3B-C). A strong increase in VGluT1 expression, detected by IF, was also observed in PTX- and Noc-treated primary neuronal cultures (Figure 3D). In order to ask whether the increased expression of VGluT1 has a functional counterpart linked to glutamatergic transmission, patch-clamp recordings of spontaneous synaptic transmission (miniature Excitatory Postsynaptic Currents, mEPSCs) on primary neuronal cultures has been performed. Consistent with our hypotheses we found a significant increase in mEPSC frequency after either PTX or Noc treatment (Figure 3E). In addition, both PTX and, albeit to a lesser extent, Noc also increased mEPSC amplitude (Figure 3E and Figure S2D).

Altogether, these results demonstrate that the modulation of the endogenous soluble Tau levels alters VGluT1 expression, ultimately impinging on glutamatergic synaptic transmission.

### VGluT1 expression is not increased by P301L mutated Tau

To investigate whether pathologically mutated Tau could have an impact on its role in the nuclear compartment, we expressed Tau^P^301^L^ and we observed that this mutant is efficiently translocated into nucleus as well as the wild type Tau. Indeed, it is detectable in both the soluble and chromatin-bound fractions (Figure 4A). We observed that Tau^P^301^L^ does not increase VGluT1 expression with respect to control cells. Indeed, both the protein (1.16±0.13SE FC) and mRNA levels (1.04±0.23SE FC) are not statistically different from those in control cells (Figure 4). Taken together, these results indicate that P301L mutation does not affect the nuclear translocation but abolishes the ability of Tau to induce VGluT1 expression and thus leads to a loss of this nuclear function.

**Figure 4.**
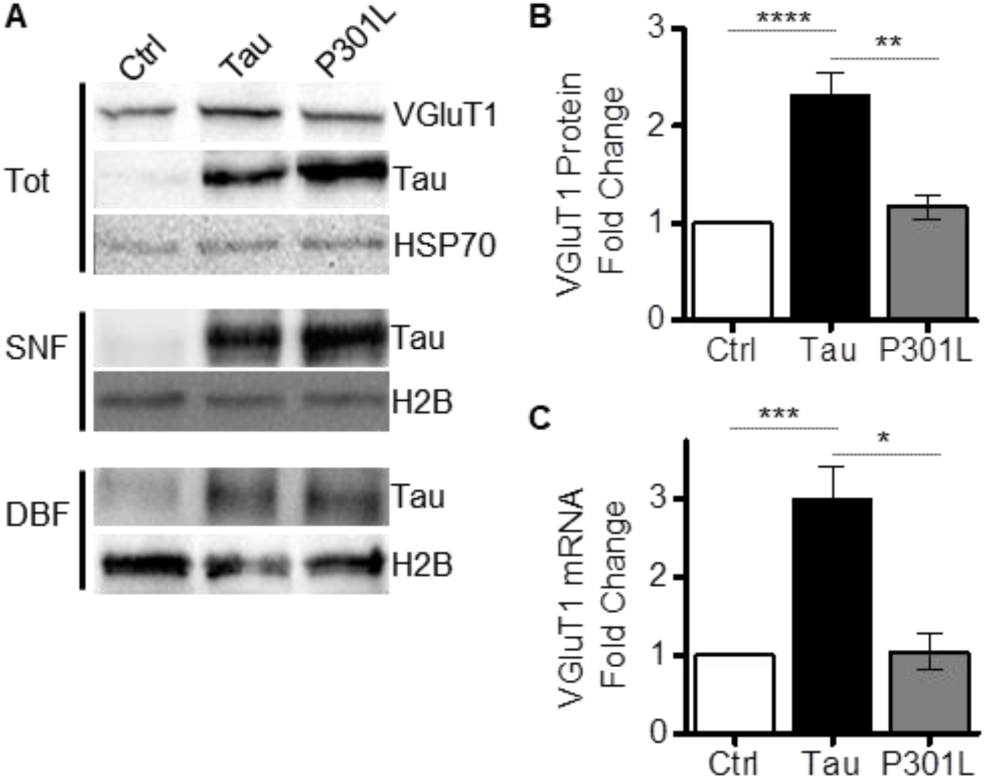
P301L mutation leads to loss of nuclear function A. Western blot of subcellular fractions obtained from HT22 cells expressing wild-type Tau or Tau^P301L^. **B**. Quantification of WB experiments. **C**. VGluT1 mRNA quantification by qPCR in HT22 cells expressing wild-type Tau or Tau^P301L^ (Mann-Whitney test; **** p<0.0001; *** p<0.001; n.s. p>0.05) (See also Figure S4).

## Discussion

Here we demonstrated that Tau, a well-established microtubule-associated protein [1,2,4,35], has a relevant role in modulating the expression of a gene specifically involved in glutamatergic synaptic transmission. We found that this function is specifically exerted by Tau protein localized in the nuclear compartment and that it is abolished by the tauopathy mutation Tau^P^301^L^. Previous evidence identified Tau in the nuclei of neuroblastoma and neuronal cells [9,13,20]. We detected Tau in the nuclei of HeLa and HT22 cells confirming that the ability of Tau to translocate into the nucleus is retained also in non-neuronal cell lines. The mechanisms by which Tau localizes to the nucleus are still unknown, even though it has been shown that the interaction of Tau with TRIM28 stabilizes and promotes Tau nuclear accumulation. In addition, both nuclear Tau and nuclear TRIM28 are increased in AD brains [36].

We observed that, in the nuclear compartment, Tau is detectable in the soluble nuclear fraction and in the DNA-bound fraction. Our demonstration of Tau association to chromatin confirms, in a cellular context, previous *in vitro* studies demonstrating a possible interaction of Tau with DNA [17,27,37].

Up to now, the role of Tau in the nucleus is still unclear. Previous studies have suggested its involvement in nucleolar organization and in DNA protection under stress conditions [14,16]. Nuclear Tau has been found to accumulate in heterochromatin during aging [21] suggesting a role in modulating gene expression but no evidence for this role is available. We focused on this hypothesis, and, in particular, on the possibility that nuclear Tau might modulate the expression of genes involved in glutamatergic synaptic transmission. Indeed, a common event identified during the progression of Tau pathology is a strong change in synaptic transmission. In particular, in the early pathologic stage, glutamate release is increased and correlates with toxic circuit hyperexcitability; in subsequent stages, this is followed by a glutamate release impairment [25,26,38–40]. Synaptic glutamate release is mediated by several proteins and the VGluT family has a relevant role by mediating glutamate loading into synaptic vesicles [28]. In tauopathy mouse models, glutamate release and VGluT1 levels are increased in the pre-symptomatic phase of the disease [25,26]. A glutamate reduction is observed in late stages of tauopathy [26,38], suggesting that alterations of VGluT1 expression are dependent on both the stage of disease and the state of Tau protein. Thus, in early phase of the disease, Tau is detached from MTs but found mostly in a soluble state while, in later stages, it is in an aggregated state. It is conceivable that the nuclear role of Tau in these two stages is distinct. We found that increasing the expression of Tau leads to a selective increase in VGluT1 expression, but not of VGLuT2 nor of VGluT3 (data not shown). Remarkably, forcing the nuclear localization of Tau (Tau-NLS) increases VGluT1 expression at the same level attained by overexpression of untagged Tau. This might indicate either that the amount of untagged Tau reaching the nucleus is saturating, or that nuclear Tau is not responsible for that increase. We clarified this point by forcing the export of Tau from the nucleus (Tau-NES). Remarkably, we found that, in this case, the expression of VGluT1 returns to control level. We conclude that the modulation of VGluT1 expression is specifically triggered by nuclear Tau.

To provide further supporting evidence, we treated cells with PTX and Noc, chemicals that increase the Tau soluble pool by different mechanisms (Di Primio et al., 2017; Kar et al., 2003). Both treatments increase Tau in the nucleus, highlighting an equilibrium between cytoplasmic and nuclear Tau. Concomitantly, the expression of VGluT1 mRNA and protein was increased. This demonstrates that Tau detachment from MTs not only affects MTs stability, but also favours its nuclear translocation, thus directly affecting VGluT1 expression. Remarkably, the increased nuclear Tau and VGluT1 expression was accompanied by an increased spontaneous synaptic transmission. In particular, the higher mEPSC frequency indicates a presynaptic effect that can be directly linked to higher VGluT1 levels. This electrophysiological observation represents an *in vitro* correlation of the neuronal hyperexcitability, a hallmark of early-stage tauopathy [25,26].

We also observed an increase of mEPSC amplitude following PTX and Noc treatments (Figure S3), which is usually attributed to a post-synaptic effect, suggesting that nuclear Tau might likewise influence the expression of glutamate receptors or of glutamate receptor interacting proteins. The modulation of VGluT1 mRNA by Tau might be exerted at the transcriptional level and Tau possible interaction with regulatory elements, i.e., enhancers or silencers might not be excluded. Previous observations clearly suggested the involvement of Tau on chromatin remodelling in pathology [41], but this was not directly linked to nuclear Tau functions. In addition, it has been recently observed that Tau activates transposable elements in Alzheimer’s disease [42].

A post-transcriptional effect of nuclear Tau on VGluT1 mRNA stability is another possibility that would lead to increased levels. In addition, an effect of nuclear Tau in promoting the export of VGluT1 mRNA to the cytoplasm could be present. It also remains to be verified whether Tau modulates the expression of other disease-relevant genes.

This nuclear function of wild type Tau might have a relevant role in the onset of the sporadic pathology as it has been widely demonstrated that the glutamate release is increased and causes hippocampal hyperexcitability in the pre-symptomatic phase of tauopathies [26,38–40]. To investigate whether mutations of Tau found in human tauopathies influence the ability to modulate gene expression, we measured VGluT1 expression in cells expressing Tau^P^301^L^, a well-characterized mutant related to a group of genetic tauopathies [3,8,43,44]. Intriguingly, Tau^P^301^L^ nuclear translocation was not affected but its overexpression had no significant effect on VGluT1 level, thus indicating a loss of nuclear function. This result could be due to a failure in the interaction of mutated Tau with a nuclear cofactor involved in gene expression. We previously demonstrated that P301L mutation induces an altered protein conformation in the cytoplasm [31]. We cannot exclude that this alteration could contribute in hindering the binding to a cofactor. Moreover, the P301L mutation is placed in the second repeat of the microtubule-binding domain (MTBD), that seems to be also crucial for DNA binding [17], therefore an altered conformation of the repeat could impact on DNA binding and, consequently, on gene expression. Although we cannot explain the molecular mechanisms involved, a more extensive study on gene expression might be performed to elucidate the role of wild type and mutated Tau on the modulation of genes involved in glutamatergic synaptic transmission.

Altogether, these results highlight the role of nuclear Tau in gene expression. This function could have a potential therapeutic relevance, and screening assays based on targeting this novel physiological mechanism will form the basis for new drug discovery approaches to fight neurodegeneration. To this regard, several aspects are still unclear, in particular, the mechanisms involved in this function, which will be the target of future research efforts. It is currently accepted that, in tauopathies and in AD, Tau gains a toxic function, linked to its detachment from MTs and to the acquisition of aggregating properties. Our results open the intriguing possibility that in early phases of neuron degeneration progression, the same process that leads to Tau detachment from MTs might trigger an additional, but not necessarily alternative, gain of function, linked to modulation of pathologically relevant genes. As the neurodegeneration progresses, the increased pool of soluble Tau starts aggregating leading to a loss of the nuclear function of Tau. This would explain the observed reduction of glutamatergic transmission and VGluT1 expression in tauopathy models and human brains. These results prompt investigations to evaluate nuclear Tau as an attractive new therapeutic target.

## SUPPLEMENTAL INFORMATION

**Figure S1.**
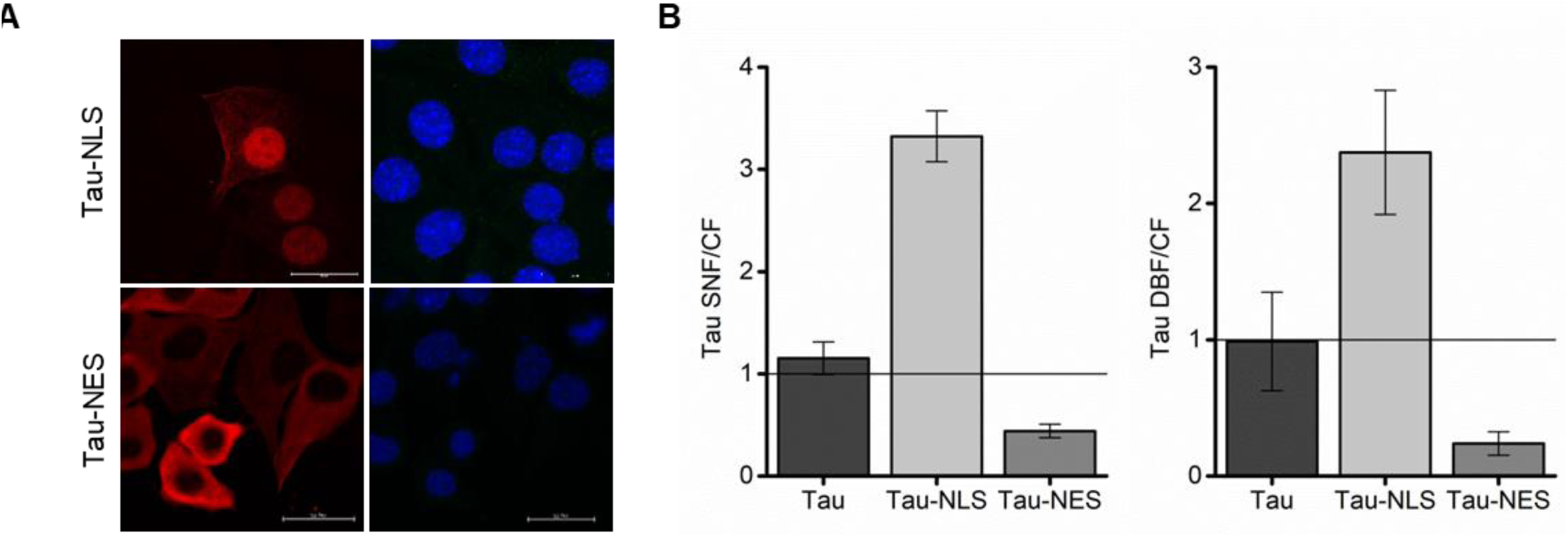
**A.** IF of HT22 cells expressing Tau-NLS or Tau-NES constructs. Tau (red), DAPI (blue). **B**. Quantification of WB in Fig.2. The ratio SNF/CF and DBF/CF=1 corresponds to Tau ratio in control cells.

**Figure S2.**
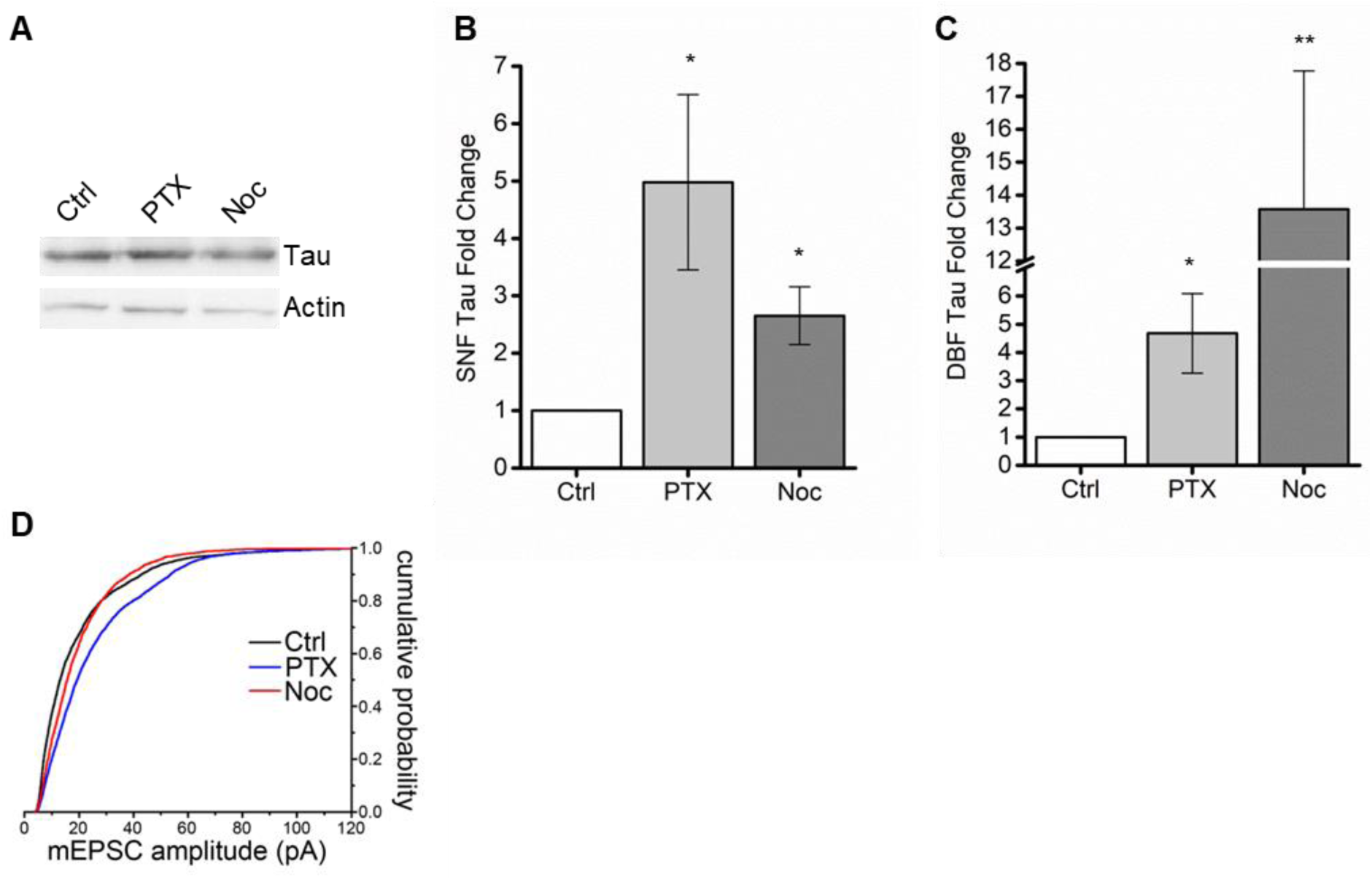
**A.** Western blot analysis of total extracts from untreated and treated HT22 cells. UT (untreated), PTX (paclitaxel), Noc (nocodazole). **B.** Quantification of Tau protein level in the soluble nuclear fraction (Mann-Whitney test; * p-value<0.05) **C.** Quantification of Tau protein level in the chromatin-bound fraction (Mann-Whitney test; ** p-value<0,01). **D.** Cumulative distributions showing mEPSC amplitude following either PTX or Noc treatment, compared to untreated controls (Ctrl, n = 3015 events from 12 cells; PTX, n = 4921 events from 10 cells; Noc, n = 3582 events from 6 cells; Kolmogorov-Smirnov test, Ctrl vs. PTX, p < 0.001; Ctrl vs. Noc, p < 0.001; PTX vs. Noc, p < 0.001).

## Experimental Procedures Chimeric Constructs Cloning

The cDNA encoding Tau isoform D (383aa) has been cloned into the BglII site of pcDNA3.1. Tau-NLS has been generated by digestion and cloning of the 3xNLS in the XhoI/BamHI site. Tau-NES has been obtained by PCR amplification of NES and cloning in EcoRI/BamHI site.

## Cell Culture, transfections and treatments

HeLa cells and immortalized hippocampal neurons HT22 were maintained in Dulbecco’s modified Eagle’s medium (DMEM) (GIBCO) supplemented with 10% FBS. The day before the experiment cells were seeded at 10^5^ cells in six-well plates or in Willco dishes (Willcowells). Lipofection was carried out with Effectene (QIAGEN) or Lipofectamine 2000 (Thermo-Fisher) according to manufacturer’s instructions.

Primary hippocampal neurons were obtained from postnatal day (P) 0 B6/129 mice. Hippocampi were dissected and triturated in cold calcium-free Hank’s balanced salt solution with 100 U ml^-1^ penicillin, 0.1 mg ml^-1^ streptomycin, and digested in 0.1% trypsin, followed by inactivation in 10% FBS DMEM (GIBCO) with 100 U ml^-1^ DNase. Neurons were seeded on poly-D-lysine-coated glass coverslips. For initial plating, neurons were maintained in Neurobasal-A medium (Invitrogen) supplemented with 4.5 g l^-1^ D-glucose, 10% FBS, 2% B27 (Invitrogen), 1% Glutamax (Invitrogen), 1 mM pyruvate, 4 μM reduced glutathione, and 12.5 μM glutamate. From the following day on, neurons were grown in Neurobasal-A medium (Invitrogen) supplemented with 2% B27 (Invitrogen), 1% Glutamax (Invitrogen), and 1 μg ml^-1^ gentamicin. For IF experiments, neurons at day *in vitro* (DIV) 18 have been used. HT22 cells and primary hippocampal neurons were treated with PTX 1µM or Noc 1µM for 4 hours.

## Western Blot and Immunostaining

Total protein extracts were prepared in lysis buffer supplemented with protease and phosphatase inhibitors. The Subcellular Protein Fractionation Kit for Cultured Cells (Thermo-Fisher) was used according to manufacturer’s instructions. For each sample 20 μg of each fraction were loaded. Proteins were separated by SDS-PAGE and electro-blotted onto Hybond-C-Extra (Amersham Biosciences) nitrocellulose membranes. Membranes were blocked (5% skimmed milk powder in TBS, 0.1% Tween 20).

For IF experiments, cells were fixed with ice-cold 100% methanol for 5 min. After permeabilization (PBS, 0.1% Triton-X100) samples were blocked (1% wt/vol BSA) and incubated with primary and secondary antibodies. Slides were mounted with Vectashield mounting medium (Vector Laboratories).

Primary antibodies for WB: mouse anti-tau (Tau5) 1:1000 ab80579 (AbCam); mouse anti-α-Tubulin Clone B-5-1-2 1:10000 (SIGMA-ALDRICH); mouse anti-GAPDH 1:15000 (Fitzgerald); rabbit anti-VGluT1 ab77822 1:500 (AbCam). Secondary antibodies for Western blot analysis were HRP-conjugated anti-mouse or anti-rabbit, purchased from Santa Cruz Biotechnology, Inc., Santa Cruz, CA, USA.

Primary antibodies for IF: mouse anti-tau (Tau-13) 1:500 (Santa Cruz); rabbit anti-VGLUT1 ab77822 1:500 (AbCam); DAPI 1:15000 (Sigma). Secondary antibodies for IF: Alexa Fluor 633; Alexa Fluor 488 (Life Technologies).

## Real-time PCR

Total RNA was extracted by Nucleospin (Macherey-Nagel) and retro-transcribed by Reverse Transcriptase Core kit (Eurogentec) according to manufacturer’s instructions. Real-time PCR was performed using the iTaqTM Universal SYBR^®^ Green Supermix (BioRad), and performed for 40 cycles of amplification with denaturation at 95°C for 15 s, annealing at 60°C for 25 s, extension at 72°C for 20 s. The primers employed were: Actin: fwd 5’-TCCATCCTGGCCTCACTGTCCAC-3’, rev 5’-GAGGGGCCGGACTCATCGTACT-3’;Tau:fwd5’-GTGACCTCCAAGTGTGGCTCATT-3’,rev5’-CTTCGACTGGACTCTGTCCTTG-3’; VGluT1: fwd 5’-GAGGAGTGGCAGTACGTGTTCC-3’, rev 5’-TCTCCAGAAGCAAAGACCCC-3’.

## Image Acquisition and Analysis

A Leica TCS SP8 confocal laser-scanning microscope (Leica Microsystems, Mannheim, Germany) equipped with Leica Application Suite (LAS) X software was used. All frames were captured by means of HC PL APO CS2 40X/1.30 (*) oil objective, a format size of 2048 × 2048 pixel and a sequential scan procedure. All confocal frames were taken by a suitable scanning power and speed along with gain level to achieve the greater signal definition and avoid any background noise. (*) N.A.=1.30.

## Patch-clamp recordings and electrophysiological data analysis

Recordings were performed on primary neuronal cultures, by adapting the procedure described in Piacentini et al. [45]. Cells were continuously bathed using Tyrode’s solution containing (in mM): NaCl 150, KCl 4, MgCl_2_ 1, CaCl2 4, Glucose 10, HEPES 10, pH 7.4 with NaOH. Borosilicate glass pipettes were pulled with a P-97 puller (Sutter, CA) to a resistance of 5-6 MΩ when filled with an internal solution containing (in mM): K-Gluconate 145, MgCl2 2, HEPES 10, EGTA 0.1, Mg-ATP 2.5, Na-GTP 0.25, phosphocreatine 5, pH 7.35 with KOH. Miniature Excitatory Postsynaptic Currents (mEPSCs) were recorded while holding the neuron at a command potential of −70 mV. Data were acquired using a MultiClamp 700A amplifier, connected to a Digidata 1322A digitizer (Molecular Devices, CA). Data were analyzed using Clampfit 10.7 (Molecular Devices) as described in Mainardi et al. [46]. An event template was constructed and used to detect mEPSCs; stringency of detection was ensured by setting the “template match threshold” to 4. After event detection, the cumulative distributions for mEPSC amplitude and frequency (the latter as interevent interval) were calculated.

## Statistical analysis

For Western Blot and quantitative real-time PCR, statistical significance was assessed by non-parametric Kruskal-Wallis test followed by pairwise Mann-Whitney test (one tailed). For qPCR, gene expression was calculated with Pfaffl method [47]. Each sample was run in triplicate and at least three biological replicates were performed for each experiment. All results are shown as mean±SEM from at least three independent experiments. Significance is indicated as * for p<0.05, ** for p<0.01, *** for p<0.001 and **** for p<0.0001. For patch-clamp recordings, statistical significance was assessed using the Kolmogorov-Smirnov test.

## Author Contributions

Conceptualization, G.S., C.D.P., V.Q. A.C.; Methodology, G.S., C.D.P., V.Q.; Investigation, G.S., M.V., M.C.C., M.M., C.I.; Writing – Original Draft, G.S, C.D.P.; Writing –Review & Editing, G.S, C.D.P., V.Q., A.C.; Funding Acquisition, C.D.P. and A.C.

## Acknowledgments

This work was supported by grants from Scuola Normale Superiore (SNS14_B_DIPRIMIO; SNS16_B_DIPRIMIO). The authors are grateful to F. Gobbo for the technical support in primary culture, A. Cellerino, F. Cremisi, M. Costa for valuable discussions.

## References

1. Arendt T, Stieler JT, Holzer M (2016) Tau and tauopathies. Brain Res Bull 126: 238–292.

2. Guo T, Noble W, Hanger DP (2017) Roles of tau protein in health and disease. Acta Neuropathol 133: 665–704.

3. Hasegawa M, Smith MJ, Goedert M (1998) Tau proteins with FTDP-17 mutations have a reduced ability to promote microtubule assembly. FEBS Lett 437: 207–210.

4. DiTella M, Feiguin F, Morfini G, C??ceres A (1994) Microfilament???associated growth cone component depends upon Tau for its intracellular localization. Cell Motil Cytoskeleton 29: 117–130.

5. Nam W, Epureanu BI (2017) Dynamic model for kinesin-mediated long-range transport and its local traffic jam caused by tau proteins. Phys Rev E 95: 1–11.

6. Vershinin M, Carter BC, Razafsky DS, King SJ, Gross SP (2007) Multiple-motor based transport and its regulation by Tau. Proc Natl Acad Sci 104: 87–92.

7. Mackenzie IRA, Neumann M (2016) Molecular neuropathology of frontotemporal dementia: insights into disease mechanisms from postmortem studies. J Neurochem 138: 54–70.

8. S. Mirra S, Murrell J, Gearing M, Spillantini MG, Goedert M, Anthony Crowther R, Levey A, Jones R, Green J, Shoffner J, et al. (1999) Tau Pathology in a Family with Dementia and a P301L Mutation in Tau. J Neuropathol Exp Neurol 58:.

9. Greenwood JA, Johnson GVW (1995) Localization and in Situ Phosphorylation State of Nuclear Tau. Exp Cell Res 220: 332–337.

10. Hua Q, He RQ (2003) Tau could protect DNA double helix structure. Biochim Biophys Acta - Proteins Proteomics 1645: 205–211.

11. Jadhav S, Cubinkova V, Zimova I, Brezovakova V, Madari A, Cigankova V, Zilka N (2015) Tau-mediated synaptic damage in Alzheimer’s disease. Transl Neurosci 6: 214–226.

12. Manczak M, Reddy PH (2012) Abnormal interaction of VDAC1 with amyloid beta and phosphorylated tau causes mitochondrial dysfunction in Alzheimer’s disease. Hum Mol Genet 21: 5131–5146.

13. Rady RM, Zinkowski RP, Binder LI (1995) Presence of tau in isolated nuclei from human brain. Neurobiol Aging 16: 479–486.

14. Sjoberg MK (2006) Tau protein binds to pericentromeric DNA: a putative role for nuclear tau in nucleolar organization. J Cell Sci 119: 2025–2034.

15. Sotiropoulos I, Galas M-C, Silva JM, Skoulakis E, Wegmann S, Maina MB, Blum D, Sayas CL, Mandelkow E-M, Mandelkow E, et al. (2017) Atypical, non-standard functions of the microtubule associated Tau protein. Acta Neuropathol Commun 5: 91.

16. Violet M, Delattre L, Tardivel M, Sultan A, Chauderlier A, Caillierez R, Talahari S, Nesslany F, Lefebvre B, Bonnefoy E, et al. (2014) A major role for Tau in neuronal DNA and RNA protection in vivo under physiological and hyperthermic conditions. Front Cell Neurosci 8: 1–11.

17. Qi H, Cantrelle F-X, Benhelli-Mokrani H, Smet-Nocca C, Buée L, Lippens G, Bonnefoy E, Galas M-C, Landrieu I (2015) Nuclear Magnetic Resonance Spectroscopy Characterization of Interaction of Tau with DNA and Its Regulation by Phosphorylation. Biochemistry 54: 1525–1533.

18. Wei Y, Qu MH, Wang XS, Chen L, Wang DL, Liu Y, Hua Q, He RQ (2008) Binding to the minor groove of the double-strand, Tau protein prevents DNA damage by peroxidation. PLoS One 3:.

19. Padmaraju V, Indi SS, Rao KSJ (2010) New evidences on Tau-DNA interactions and relevance to neurodegeneration. Neurochem Int 57: 51–57.

20. Loomis P a, Howard TH, Castleberry RP, Binder LI (1990) Identification of nuclear tau isoforms in human neuroblastoma cells. Proc Natl Acad Sci U S A 87: 8422–8426.

21. Gil L, Federico C, Pinedo F, Bruno F, Rebolledo AB, Montoya JJ, Olazabal IM, Ferrer I, Saccone S (2017) Aging dependent effect of nuclear tau. Brain Res 1677: 129–137.

22. Rossi G, Dalprà L, Crosti F, Lissoni S, Sciacca FL, Catania M, Fede D, Mangieri M, Giaccone G, Croci D, et al. (2008) A new function of microtubule-associated protein tau. Involvement in chromosome stability. Cell Cycle 7: 1788–1794.

23. Rossi G, Conconi D, Panzeri E, Redaelli S, Piccoli E, Paoletta L, Dalprà L, Tagliavini F (2013) Mutations in MAPT gene cause chromosome instability and introduce copy number variations widely in the genome. J Alzheimer’s Dis 33: 969–982.

24. Rossi G, Conconi D, Panzeri E, Paoletta L, Piccoli E, Ferretti MG, Mangieri M, Ruggerone M, Dalprà L, Tagliavini F (2014) Mutations in MAPT give rise to aneuploidy in animal models of tauopathy. Neurogenetics 15: 31–40.

25. Hunsberger HC, Rudy CC, Batten SR, Gerhardt GA, Reed N (2015) P301L Tau Expression Affects Glutamate Release and Clearance in the Hippocampal Trisynaptic Pathway. J Neurochem 132: 169–182.

26. Crescenzi R, DeBrosse C, Nanga RPR, Byrne MD, Krishnamoorthy G, D’Aquilla K, Nath H, Morales KH, Iba M, Hariharan H, et al. (2017) Longitudinal imaging reveals sub-hippocampal dynamics in glutamate levels associated with histopathologic events in a mouse model of tauopathy and healthy mice. Hippocampus 27: 285–302.

27. Krylova SM, Musheev M, Nutiu R, Li Y, Lee G, Krylov SN (2005) Tau protein binds single-stranded DNA sequence specifically - The proof obtained in vitro with non-equilibrium capillary electrophoresis of equilibrium mixtures. FEBS Lett 579: 1371–1375.

28. Kaneko T, Fujiyama F (2002) Complementary distribution of vesicular glutamate transporters in the central nervous system. Neurosci Res 42: 243–250.

29. Alonso A del, Li B, Grundke-Iqbal I, Iqbal K (2008) Mechanism of Tau-Induced Neurodegeneration in Alzheimer Disease and Related Tauopathies. Curr Alzheimer Res 5: 375–384.

30. Iqbal K, Liu F, Gong C-X, Grundke-Iqbal I (2010) Tau in Alzheimer Disease and Related Tauopathies. Curr Alzheimer Res 7: 656–664.

31. Di Primio C, Quercioli V, Siano G, Rovere M, Kovacech B, Novak M, Cattaneo A (2017) The Distance between N and C Termini of Tau and of FTDP-17 Mutants Is Modulated by Microtubule Interactions in Living Cells. Front Mol Neurosci 10: 1–13.

32. Jordan M a, Thrower D, Wilson L (1992) Effects of vinblastine, podophyllotoxin and nocodazole on mitotic spindles. Implications for the role of microtubule dynamics in mitosis. J Cell Sci 102 (Pt 3: 401–416.

33. Breuzard G, Hubert P, Nouar R, De Bessa T, Devred F, Barbier P, Sturgis JN, Peyrot V (2013) Molecular mechanisms of Tau binding to microtubules and its role in microtubule dynamics in live cells. J Cell Sci 126: 2810–2819.

34. Kar S, Fan J, Smith MJ, Goedert M, Amos LA (2003) Repeat motifs of tau bind to the insides of microtubules in the absence of taxol. EMBO J 22: 70–77.

35. Bakota L, Brandt R (2016) Tau Biology and Tau-Directed Therapies for Alzheimer’s Disease. Drugs 76: 301–313.

36. Rousseaux MWC, de Haro M, Lasagna-Reeves CA, de Maio A, Park J, Jafar-Nejad P, Al-Ramahi I, Sharma A, See L, Lu N, et al. (2016) TRIM28 regulates the nuclear accumulation and toxicity of both alpha-synuclein and tau. Elife 5: 1–24.

37. P. Vasudevaraju EG, Hegde ML, Collen TB, Britton GB, Rao KS (2012) New evidence on α-synuclein and Tau binding to conformation and sequence specific GC* rich DNA: Relevance to neurological disorders. J Pharm Bioallied Sci 4: 112–117.

38. Maurin H, Chong SA, Kraev I, Davies H, Kremer A, Seymour CM, Lechat B, Jaworski T, Borghgraef P, Devijver H, et al. (2014) Early structural and functional defects in synapses and myelinated axons in stratum lacunosum moleculare in two preclinical models for tauopathy. PLoS One 9:.

39. Dickerson BC, Salat DH, Greve DN, Chua EF, Rand-Giovannetti E, Rentz DM, Bertram L, Mullin K, Tanzi RE, Blacker D, et al. (2005) Increased hippocampal activation in mild cognitive impairment compared to normal aging and AD. Neurology 65: 404–411.

40. Reiman EM, Quiroz YT, Fleisher AS, Chen K, Velez-Pardo C, Jimenez-Del-Rio M, Fagan AM, Shah AR, Alvarez S, Arbelaez A, et al. (2012) Brain imaging and fluid biomarker analysis in young adults at genetic risk for autosomal dominant Alzheimer’s disease in the presenilin 1 E280A kindred: a case-control study. Lancet Neurol 11: 1048–1056.

41. Frost B, Hemberg M, Lewis J, Feany MB (2014) Tau promotes neurodegeneration through global chromatin relaxation. Nat Neurosci 17: 357–366.

42. Guo C, Jeong HH, Hsieh YC, Klein HU, Bennett DA, De Jager PL, Liu Z, Shulman JM (2018) Tau Activates Transposable Elements in Alzheimer’s Disease. Cell Rep 23: 2874–2880.

43. Hutton M, Lendon CL, Rizzu P, Baker M, Froelich S, Houlden HH, Pickering-Brown S, Chakraverty S, Isaacs A, Grover A, et al. (1998) Association of missense and 5’-splice-site mutations in tau with the inherited dementia FTDP-17. Nature 393: 702–704.

44. Fischer D, Mukrasch MD, Von Bergen M, Klos-Witkowska A, Biemat J, Griesinger C, Mandelkow E, Zweckstetter M (2007) Structural and microtubule binding properties of tau mutants of frontotemporal dementias. Biochemistry 46: 2574–2582.

45. Piacentini R, Li Puma DD, Mainardi M, Lazzarino G, Tavazzi B, Arancio O, Grassi C (2017) Reduced gliotransmitter release from astrocytes mediates tau-induced synaptic dysfunction in cultured hippocampal neurons. Glia 65: 1302–1316.

46. Mainardi M, Spinelli M, Scala F, Mattera A, Fusco S, D’Ascenzo M, Grassi C (2017) Loss of Leptin-Induced Modulation of Hippocampal Synaptic Trasmission and Signal Transduction in High-Fat Diet-Fed Mice. Front Cell Neurosci 11: 1–11.

47. Pfaffl MW (2001) A new mathematical model for relative quantification in real-time RT-PCR. Nucleic Acids Res 29: 45e–45.

